# Stereotypical hippocampal clustering predicts navigational success in virtualized real-world environments

**DOI:** 10.1101/2023.03.23.533994

**Authors:** Jason D. Ozubko, Madelyn Campbell, Abigail Verhayden, Brooke Demetri, Molly Brady, Iva Brunec

**Affiliations:** Psychology Department SUNY Geneseo; Psychology Department; University of Pennsylvania & Temple University

**Author notes:** Corresponding Author: Jason D. Ozubko. Author Contributions JDO was responsible for the conceptualization, design, analyses, supervision, and writing. MC, AV, BD, and MB all were involved in data collection and curation, investigation, and project administration. MB and BD also assisted with design and supervision. IB assisted with conceptualization and writing.

## Abstract

Structural differences along the long-axis of the hippocampus have long been believed to underlie meaningful functional differences, such as the granularity of information processing. Recent findings show that data-driven parcellations of the hippocampus sub-divide the hippocampus into a 10-cluster map with anterior-medial, anterior-lateral, and posteroanterior-lateral, middle, and posterior components. We tested whether task and experience could modulate this clustering using a spatial learning experiment where subjects were trained to virtually navigate a novel neighborhood in a Google Street View-like environment over a two-week period. Subjects were scanned while navigating routes early in training and at the end of their two-week training. Using the 10-cluster map as the ideal template, we find that subjects who eventually learn the neighborhood well have hippocampal cluster-maps consistent with the ideal—even on their second day of learning—and their cluster mappings do not change over the two week training period. However, subjects who eventually learn the neighborhood poorly begin with hippocampal cluster-maps inconsistent with the ideal, though their cluster mappings become more stereotypical by the end of the two week training. Interestingly this improvement seems to be route specific as even after some early improvement, when a new route is navigated participants’ hippocampal maps revert back to less stereotypical organization. We conclude that hippocampal clustering is not dependent solely on anatomical structure, and instead is driven by a combination of anatomy, task, and importantly, experience. Nonetheless, while hippocampal clustering can change with experience, efficient navigation depends on functional hippocampal activity clustering in a stereotypical manner, highlighting optimal divisions of processing along the hippocampal anterior-posterior and medial-lateral-axes.

## Main

Structural differences along the long-axis of the hippocampus have long been believed to underlie meaningful functional differences. For instance, the granularity of information processing is believed to vary along the hippocampus, with coarse- to fine-grained processing roughly mapping on to the anterior and posterior regions respectively (1–4). These conclusions are drawn from functional analyses that have often segmented the hippocampus using anatomical borders along the long-axis, with the assumption that boundaries in structure define boundaries in function. Recently, however, data-driven parcellation of the hippocampus using group masked independent components analysis (mICA) has shown that the bilateral hippocampi cluster into ten distinct regions, five in each hemisphere, which include three anterior clusters (anterior-medial, anterior-lateral, and posteroanterio-lateral), a middle cluster, and a posterior cluster (5–8) (see Figure 2A). This 10-cluster data-driven mICA parcellation of the hippocampus (which we will subsequently refer to as the mICA-10 map), has not only been observed reliably in resting state data (5–8), but the clusters themselves have been shown to underlie meaningful differences in granularity of information processing (7).

Specifically, Thorp et al. (7) recently used the mICA-10 map to investigate inconsistencies in the literature regarding hippocampal granularity and the long-axis. In their study, Thorp et al. used the mean correlation of voxelwise timecourses or “inter-voxel similarity” (IVS) (2), to provide a proxy measure of signal complexity, and examined the impact of different parcellations of the hippocampus on IVS. Based on past research regarding granularity, IVS would be expected to decline from the anterior-to-posterior hippocampus. While traditional anterior-posterior hippocampal segmentation did find this result, ternary (anterior-middle-posterior) segmentation indicated that IVS declined in the middle of the hippocampus but not the anterior or posterior regions, and senary segmentation (segmenting the hippocampus into 6 regions along its long axis) indicated that IVS *increased* along the long-axis. Thorp et al. argued that these inconsistencies represented a significant problem in the literature, as the selection of hippocampal segmentation is often seen as relatively inconsequential and varies between studies yet carries significant implications. To tackle the issue of reproducibility then, Thorp et al. proposed that a functionally informed approach was needed for segmentation. Performing mICA on their functional data revealed the same mICA-10 map that had been found by past researchers (5–8) and more importantly, examining IVS using the mICA-10 map Thorp et al. discovered that granularity varies along the medial-lateral dimension in the anterior hippocampus and along the anterior-posterior dimension along the long-axis. These results demonstrate just how critical the segmentation of the hippocampus is, and how significantly results can change based on choice of segmentations.

Thorp et al.’s (7) work was seminal in highlighting the importance of functional parcellation. It might be tempting to view the mICA-10 map as a new standard by which to parcellate the hippocampus but how universal is the mICA-10 map? In all studies to date the mICA-10 map has only been observed from resting-state data (5–8), would the mICA-10 map replicate if the hippocampus were engaged in a task-dependent manner? By comparison, during resting state the default mode network is a functional network known to emerge but this network is not characteristic of how these brain regions regularly interact in all scenarios (9,10). Another related question is whether the mICA-10 map is emerging consistently due to the intrinsic organization of the hippocampus or whether behavioural factors like experience and performance on task could alter the hippocampal maps produced by mICA.

To further understand functional parcellations of the hippocampus, we present the first test that we are aware of to examine how task and experience can modulate functional hippocampal clustering using a spatial learning experiment. Spatial navigation was selected due to its long-standing connection with hippocampus when navigating in novel environments (11–18) plus its waning influence during navigation of well-learned environments (16,19–23). In our experiment, subjects were trained to virtually navigate a novel neighborhood in a Google Street View-like environment (Figures 1A, 1C-1E) over a two-week period (7 sessions). Subjects were scanned while navigating routes early in training (on Session 2) and at the end of their two-week training (on Sessions 6 and 7) (see Figure 1B for task design). Navigation also varied between passive (Sessions 1-5 and 7) and active (Session 6). In all cases we will compare the mICA hippocampal clusters observed in our data with the mICA-10 cluster map. Our paradigm thus allows us to explore how hippocampus parcellations may change (compared to the mICA-10 map) in novel spatial learning conditions, whether parcellation is influenced by navigational ability, whether it changes over time, and whether it changes in passive vs. active navigation conditions.

**Figure 1.**
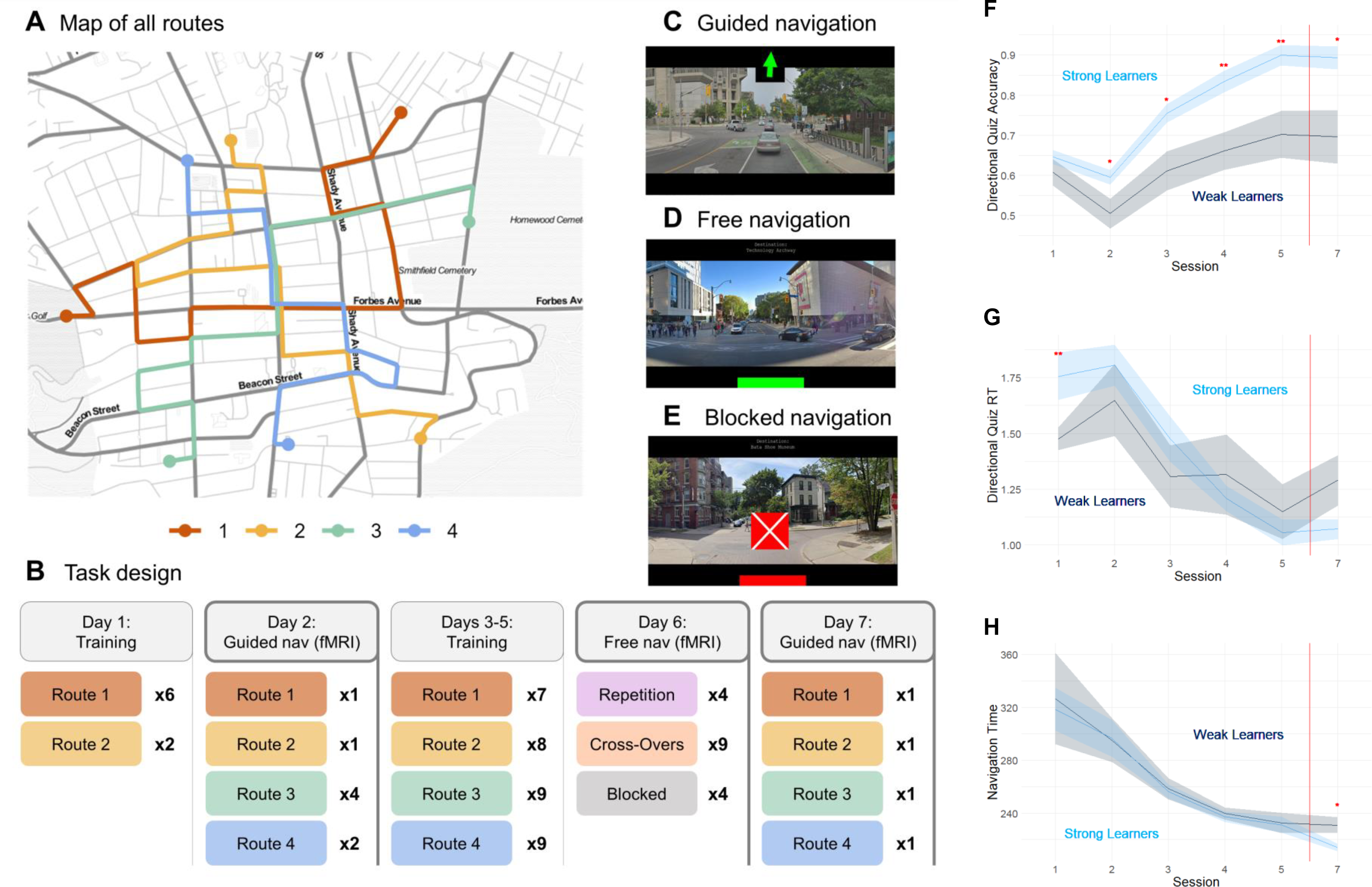
(A) An overhead view of the neighborhood that was learned in this study. The four training routes are shown, each in a separate color. During the guided training and guided navigation days, subjects were guided along the routes, having been instructed to learn the routes during this guided navigation. Each route was practiced at least 11 times over the 2-weeks of training. (C) Summary of the experimental schedule that occurred over a 2-week period. On Day 2, subjects were scanned to get an early measure of neural activity during guided navigation. Subjects were also scanned on Day 7 to get a late measure of neural activity during guided navigation. On Day 6, subjects were presented with a series of navigational challenges included Repetition routes (navigating learned routes from memory), Cross-Over routes (starting at the beginning of one route and having to navigate to the end point of a different route), and Blocked routes (navigating along a learned route from memory except that the center of town was blocked off and had to be navigated around). Scanning on Day 6 was meant to provide a measure of neural activity during free navigation of a recently learned environment. (C) A sample depicting the subject’s first person perspective during this experiment. Subjects never saw an overhead map and did all navigation from the first-person perspective. On training and guided navigation days, after navigating a route, subjects were given a recognition and directional test to quiz the information they learned from the route. (D) A sample depicting the subject’s perspective on repetition and cross-over routes on the free navigation trials on Day 6. Subjects were given a destination name but no other guidance during navigation and had to navigate from memory. (E) A sample image depicting a blockage that a subject might encounter when navigating blocked routes on Day 6. (F) Mean directional quiz accuracy for Strong and Weak Learners from Sessions 1-5 and 7. Standard error is represented with the shaded areas around the line. Strong Learners showed significantly more accurate performance than Weak Learners in starting in Session 2. (G) Mean directional quiz reaction time (RT) for Strong and Weak Learners from Sessions 1-5 and 7. Standard error is represented with the shaded areas around the line. Strong Learners were significantly slower than Weak Learners in Session 1 but not at any other times. (H) Mean overall navigation time for Strong and Weak Learners from Sessions 1-5 and 7. Standard error is represented with the shaded areas around the line. Strong Learners were significantly faster than Weak Learners in Session 7 but not at any other times. Note in Figures F-H significance is indicated as * *p* < .05 and ** *p* < .01.

## Results

Although navigational ability certainly varies along a continuum, mICA analyses must be performed on discrete groups. Hence, to examine the role of navigational ability using mICA, participants were divided into Strong and Weak Learners based on their performance during the two week training period. These two groups did not differ on weekly hours spent playing video games, *t*(29) = 0.51, *p* = .61, navigation strategies, *t*(29) = 0.78, *p* = .44, or sense of direction, *t*(29) = 0.36, *p* = .72, sense of direction, *t*(29) = 0.78, *p* = .44, or age, *t*(29) = 1.00, *p* = .33.

Nonetheless, Strong Learners were significantly more accurate during navigational directional quizzes than Weak Learners, with a mean accuracy of .90 (SE = .02) after the 5^th^ session compared to a mean accuracy of .70 (SE = .04) for the Weak learners, *t*(29) = 4.48, *p* < .01 (Figure 1F). In fact, across all training sessions except for the first, Strong Learners were more accurate than Weak Learners in directional quiz accuracy, all *p*’s < .05. Interestingly, Strong Learners were slower to respond to directional quizzes than were Weak Learners during the first training session, *t*(29) = 2.37, *p* < .05, although this difference quickly subsided with further training (see Figure 1G). There was a marginal trend for Strong Learners to be faster than Weak Learners on the directional quizzes on the very last session of the experiment, *t*(29) = 1.81, *p* = .09. Strong Learners were however, significantly faster to navigate than were Weak Learners during the very last training session, *t*(29) = 2.49, *p* < .05 (see Figure 1H). There was no other difference in navigation time in any other training sessions. On the whole then, Strong Learners differentiated themselves from Weak Learners by more accurate navigational learning, with only minimal differences in speed of navigation and no differences in general navigational traits or abilities.

### Effect of Experience and Time

Group masked independent component analyses (mICA) were run separately for Strong and Weak Learners on functional data collected during the navigation trials in Sessions 2, 6 (Strong Learners only), and 7. For these initial analyses, dimensionality of 10 was chosen based on prior research showing this to be the most reliable number of components without over-parcellating the hippocampus (5–8). These analyses separated the hippocampus into 10 clusters, based on functional activity, that maximize the spatial independence between clusters and the temporal coherence within components. In Sessions 2 and 7 participants were passively navigating along routes, whereas in Session 6 participants were actively navigating those same routes on Route Following trials. There were also Route Finding trials and Blocked Routes in Session 6 (see Methods) which will be explored in the Active vs. Passive section below, for now we report only the Route Following trials.

Our first analysis examined the mICA hippocampal clusters in the Strong and Weak Learners in Session 2 (the first scan session), compared to Session 6 (free navigation scan), and Session 7 (the last scan session) (see Figure 2C). For these clusters, 50 split-half analyses were carried out per condition to examine cluster reliability, resulting in the distributions shown in Figure 2B. Weak Learners demonstrated lower levels of cluster stability than Strong Learners in both Sessions 2, *t*(98) = 15.48, *p* < .01, and 7, *t*(98) = 11.80, *p* < .01, however, cluster stability improved between Session 2 and Session 7 for Weak Learners, *t*(98) = 5.32, *p* < .01. Strong Learners actually declined in cluster stability between Session 2 and Session 7, *t*(98) = 3.27, *p* < (and between Session 6 and Session 7, *t*(98) = 2.63, *p* < .01, but not between Session 2 and 6, *t*(98) = 0.68, *p* = .50). Nonetheless, Strong Learners generally had good levels of split-half reliability (> .80) and showed consistently more stable clusters than Weak Learners.

**Figure 2.**
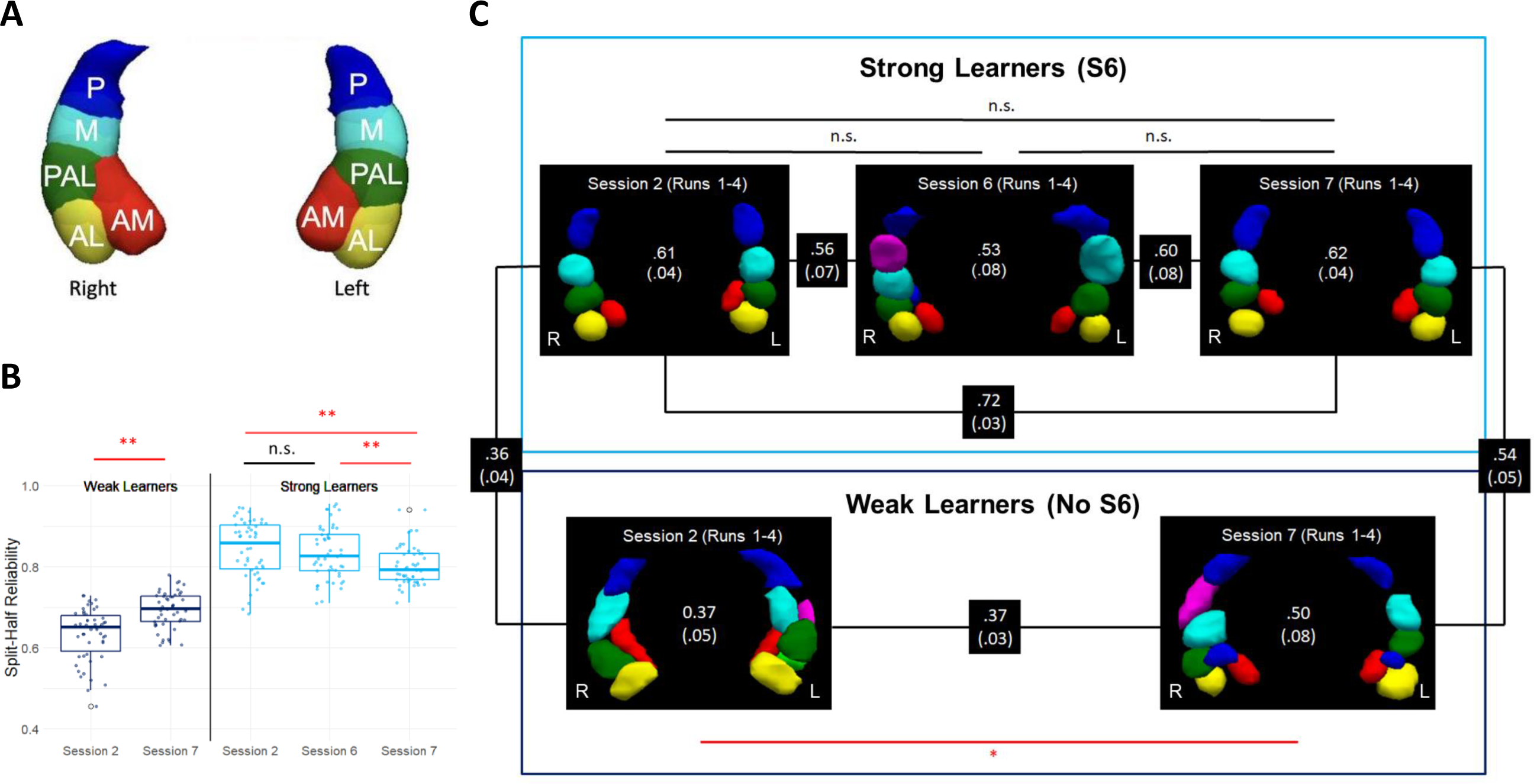
(A) The mICA-10 hippocampal clusters extracted using masked independent component analyses (mICA) from Thorp et al. (2022). In Thorp et al., mICA was applied resting state data from 183 subjects which parcellated the hippocampus into anterior-medial (AM; red), anterior-lateral (AL; yellow), posteroanterior-lateral (PAL; green), middle (M; cyan), and posterior (P; blue). These clusters were used as the ideal with which to compare the clusters extracted from our navigation study. (B) Split-half reliability distributions for strong and weak learners separated by session. These split-half analyses assess the reliability of the clusters we observed in our experiment. For these analyses, 50 split-half analyses were carried out per condition, resulting in the distributions shown. Weak learners generally demonstrated lower levels of cluster stability than strong learners, albeit cluster stability improved between Session 2 and Session 7 for weak learners. Strong learners actually declined in cluste stability at Session 7 compared to prior sessions, although overall cluster stability was always significantly higher than that of weak learners. (C) Hippocampal cluster maps derived from applying mICA to the navigation data from Session 2, Session 6, and Session 7. The statistic provided in the center of each cluster map is the mean optimized Jaccard similarity coefficient (with standard error in parentheses). This coefficient estimates the degree of fit between the observed clusters and the ideal cluster mappings as derived by Thorp et al. (1). The strong learners had cluster maps at all levels of learning and navigation that fit the idealized cluster maps fairly well (Jaccard coefficient > .5). In contrast, the poor learners had cluster maps that fit the ideal fairly poorly initially (mean Jaccard coefficient of .37) but improved over time. The significant tests highlighted in this figure refer to the difference between these Jaccard coefficients. Jaccard coefficients are also reported between cluster maps to indicate how well any two cluster maps match one another. Note that for Strong Learners in Session 6 the blue posterior cluster is bilateral; for Weak Learners in Session 2 the blue and red clusters are bilateral; and for the Weak Learners in Session 7 the blue clusters are all part of a single bilateral cluster. All other clusters within each figure are unique and separate, even if colored identically across hemispheres (e.g., the blue clusters in the left and right hemispheres are two separate clusters for all other plots, just colored the same for visualization purposes).

Jaccard similarity coefficients were used to measure the degree of fit between our observed cluster maps and the mICA-10 map. The Jaccard coefficient measures the proportion of overlap between two clusters. For our analyses, we setup a search algorithm that would compare the Jaccard coefficient between every possible pair of clusters from the mICA-10 and a target cluster map and find the pairs which together produced the highest overall fit, and the mean Jaccard coefficient was then calculated across all these pairings. This maximized mean Jaccard coefficients are reported in Figure 2C. As can be seen in the figure, Strong Learners had cluster maps at all levels of learning and navigation that fit the mICA-10 map fairly well (Jaccard coefficients > .5). Indeed, Strong Learners did not see their fit with the mICA-10 map significantly change from Session 2 to Session 6, *t*(158) = 1.32, *p* = .19, from Session 6 to Session 7, *t*(158) = 1.57, *p* = .12, or from Session 2 to Session 7, *t*(158) = 0.35, *p* = .73. In contrast, the Weak Learners had cluster maps that fit the ideal fairly poorly, with fits to the mICA-10 map being significantly worse than Strong Learners in both Session 2, *t*(122) = 5.20, *p* < .01, and Session 7, *t*(122) = 2.01, *p* < .05. However, Weak Learners did see a significant improvement in fit from Session 2 to Session 7, *t*(86) = 2.14, *p* < .05, indicating that while they began with the worst fit to the mICA-10 map, their fits were significantly improving as a result of experience.

### Novel vs. Recently Experienced Routes

In Session 2 participants navigated 4 routes: Two routes (Routes 1 and 2) had been navigated in Session 1, whereas the other two routes (Routes 3 and 4) were completely novel (see Methods). To examine the influence of recent exposure vs. novelty, we carried out separate mICA analyses on Routes 1 and 2 (recently exposed) and on Routes 3 and 4 (completely novel) in Session 2 for both Strong and Weak Learners (see Figure 3A). Strong Learners showed better fits with the mICA-10 map than the Weak Learners for novel routes, *t*(60) = 4.58, *p* < .01, but for recently introduced routes this difference was not significant, *t*(60) = 0.88, *p* = .39. Thus, prior exposure to a route had an almost immediate effect on the hippocampal parcellations for Weak Learners, however this impact was route-specific. In other words, Weak Learners were beginning to change their hippocampal parcellations almost immediately after being exposed to a route, but this change did not reflect a general shift in strategy. Instead, it seems to be the result of spatial learning along a specific route. For routes that have not yet been experienced, Weak Learners defaulted to a more disorganized hippocampal parcellation.

**Figure 3.**
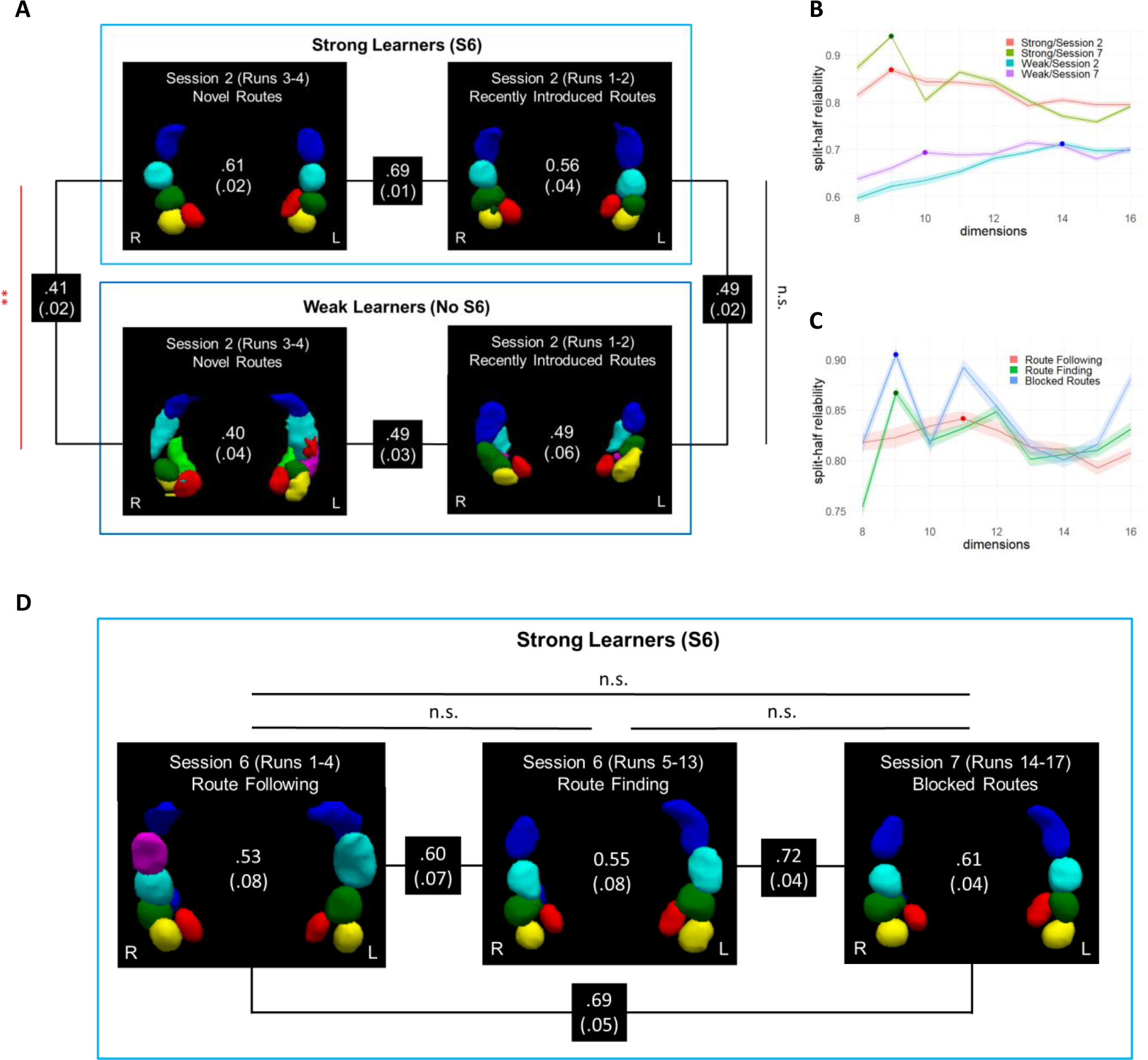
(A) Hippocampal cluster maps derived from applying mICA to novel and recently introduce routes in Session 2. Mean Jaccard similarity coefficients (and standard error) comparing each cluster map to the mICA-10 maps are reported in the center of each figure, Jaccard coefficients between each cluster map are shown between figures. (B) Dimensional reproducibility analyses of cluster maps with dimensionality ranging from 8 to 16 for Strong and Weak Learners in Sessions 2 and 7. Note that the first local maximum encountered after 8 dimensions is considered the optimal dimensionality in this case and is marked by a point for each condition. Standard error is shown in the shaded areas. (C) Dimensional reproducibility analyses of cluster maps with dimensionality ranging from 8 to 16 for Strong Learners in Route Following, Route Finding, and Blocked Routes from Session 6. The first local maximum is once again marked by a point for each condition as the optimal dimensionality. Standard error is shown in the shaded areas. (D) Hippocampal cluster maps derived from applying mICA to Route Following, Route Finding, and Blocked Routes from Session 6. Mean Jaccard similarity coefficients (and standard error) comparing each cluster map to the mICA-10 maps are reported in the center of each figure. In this case, all subjects were strong navigators. Importantly however, cluster mappings were largely consistent with the mICA-10 map and did not vary by navigation kind. The one exception is that right middle/posterior hippocampus may be more divided during Route Following compared to Route Finding and Blocked Routes. Note that for Route Following trials, the blue cluster is bilateral with an anterior portion in the left (partially obscured); and for both the Route Finding and Blocked Routes the left blue cluster also has an anterior bit (partially obscured, just under the green cluster). All other clusters within each figure are unique and separate, even if colored identically.

### Hippocampal Dimensionality

Despite Weak Learners hippocampal cluster maps improving over time, even by Session 7 they were still fitting the mICA-10 map relatively poorly. In all of our analyses thus far, we selected a dimensionality of 10 based on prior research (5–8), however, the Weak Learner’s mICA cluster maps included a number of bilateral clusters and clusters that had no obvious mICA-10 analog. Both of these features suggest the mICA analyses may have been struggling to fit a more multi-dimensional hippocampal map into a 10-dimension limit. To test this possibility, split-half reliability analyses were carried out for Sessions 2 and 7 for both Strong and Weak Learners across dimensions 8 through 16 (see Figure 3B). For these analyses, the optimal dimensionality for a condition was defined as the first local maximum encountered as dimensionality increased from 8 to 16. For Strong Learners the optimal number of dimensions was 9 in both Sessions 2 and 7. However, for Weak Learners, the optimal number of dimensions was 14 in Session 2 and 10 in Session 7. These data indicate that Weak Learners hippocampi may be over-parcellated early in training, but that over time their hippocampi reduce towards the 10 dimensions observed with the mICA-10 map. Regarding Strong Learners, they begin and remain close to the 10 dimensions observed in the past literature although interestingly, may be ideally parcellated by slightly fewer dimensions.

### Active Navigation

Up to this point analyses have focused mostly on passive navigation. In terms of active navigation, we examined more closely the hippocampal cluster maps produced at the end of training in Session 6. Because only Strong Learners completed Session 6 this analysis focuses purely on the Strong Learners. During Session 6, there were three distinct kinds of routes representing different navigational challenges: Route Following, Route Finding, and Blocked Routes (see Methods). Whereas Route Following involved following a learned route from memory, Route Finding and Blocked Routes both involved navigating through potentially unknown streets and discovering one’s own path to the goal. Though each of these routes involved different navigational challenges, applying a 10-dimensional mICA to the functional data from each revealed cluster maps that all fit the mICA-10 standard well (see Figure 3D; Jaccard coefficients > .5). Indeed, there was no significant difference between the Jaccard coefficients fitting each cluster map to the mICA-10 map between any of these conditions (all *p*’s > .33), nor was there any difference compared to the fits seen in Session 7 (passive navigation at the end of training) (all *p*’s > .15). That said, dimensionality may increase during active navigation compared to passive navigation, as on Route Following trials the middle-posterior right hippocampus seemed to sub-divide into three distinct clusters rather than two (Figure 3B). Split-half reliability analyses carried out across dimensions 8 through 16 looking for the first local maximum after 8 dimensions revealed optimal dimensions of 11 for Route Following, 9 for Route Finding, and 9 for Blocked Routes (Figure 3C). Overall then, active navigation had a minimal impact on fit compared to passive navigation, although interestingly hippocampal dimensionality may increase marginally for Route Following. Nonetheless, the 10-dimensional mICA-10 map characterized active navigation well, indicating no large scale differences from passive navigation.

## Discussion

Traditional parcellations of the hippocampus have focused on structural boundaries. Recent work, however, has demonstrated that functional parcellation of the hippocampus reveals a more complex organization of the hippocampus (5–8). To the best of our knowledge, ours is the first work to examine how hippocampal parcellation operates in task-dependent conditions and we have made several significant discoveries.

Our results show that the 10-dimensional hippocampal cluster map found in past studies (1–4) to be remarkably resilient, and that it is not limited to resting state data. Indeed, our data provides strong support for the idea that the mICA-10 map might represent an optimal organization of the hippocampus as it was those participants who performed best in our spatial navigation task who showed the clearest example of a mICA-10-like map. However, another key element of our findings is that both experience and ability yield significant influence when it comes to hippocampal parcellation: Weak Learners readily did *not* show the mICA-10 map. While Weak Learners did seem to converge towards a more mICA-10-like parcellation after 2 weeks of training, analyses revealed that their hippocampi were nonetheless still parcellating into maps that were more multi-dimensional and more distinct from the mICA-10 than Strong Learners.

Another set of key discoveries from our results is understanding how parcellations change with experience. Weak Learners did see their hippocampal parcellations become more stereotypical over time, showing that parcellations can improve with experience. Though, at present, it is difficult to say with certainty whether parcellation is a cause or a consequence of better spatial learning, our examination of Session 2 suggests that it might be a consequence: After only a few exposures to a route the day before, Weak Learners already began to show more idealized hippocampal parcellations when navigating those routes. However, when navigating novel routes they reverted back to sub-optimal hippocampal parcellations. This pattern suggests that whereas Strong Learners approach spatial navigation with more optimal hippocampal organization in place, Weak Learners only attain such organization as a consequence of learning, and this learning is route-specific.

Our data also showed that when hippocampal parcellation is sub-optimal, there is evidence that it might be over-parcellated, rather than under-parcellated. The disorganized hippocampal cluster maps of the Weak Learners often contained clusters that had no analogous cluster in the mICA-10 map and contained bilateral clusters, two indications that Weak Learners’ functional hippocampus activity might contain more than 10 components. Indeed, split-half reliability analyses which explore the impact of different levels of dimensionality found that the Weak Learners, at all levels of experience, often produced data which appeared to settle on more than 10 dimensions being the optimal fit. Interestingly, even for these more optimized fits, Weak Learners still did not produce hippocampal maps as reliable as those from Strong Learners, underscoring the fact that Weak Learners maps are inherently less stable than the more mICA-10-like maps of Strong Learners.

A final key aspect of our results is the discovery that active navigation and passive navigation show little difference in terms of parcellation. Indeed, Strong Learners actually showed little change in their cluster maps from early learning (Session 2) to later learning (Session 7) or free navigation (Session 6). Parcellation was minimally affected by whether participants were route following, route finding, or navigating around blockages. These findings make sense in the broader context of our findings, which suggest that there might be an optimal organization of the hippocampus for spatial navigation tasks, which resembled the mICA-10 map. Since Strong Learners as a group were categorized as such precisely because they were good at navigating, it is consistent to see that their hippocampal parcellations remain optimal at all stages of learning and in all kinds of navigational scenarios. Though our active navigation tasks were too challenging for Weak Learners to take part (which is partially why Weak Learners were designated as Weak Learners to begin with), it would be quite illuminating in future studies to see if Weak Learners would show unique hippocampal parcellations in active vs. passive conditions. Given that Weak Learners are less capable navigators in general, it would follow that Weak Learners might show even less idealized hippocampal parcellations during active navigation, and their hippocampal parcellations during active navigation may be distinct from their parcellations during passive navigation (unlike the Strong Learners).

When all is said and done, could the mICA-10 map represent a new standard parcellation for the hippocampus? Though we have continually argued that the mICA-10 may represent an optimal organization of the hippocampus, we have also continually reported that not all participants exhibited such a cluster map. Indeed, the blind application of the mICA-10 map to parcellate the hippocampus for further functional analysis may be ill-advised. Much like Thorp et al. (7) we would instead advocate for the importance of data-driven approaches to hippocampal parcellation, and that hippocampi may be best parcellated based on the data-at-hand for any particular study. Although the mICA-10 map may represent an optimal parcellation of the hippocampus, there is no guarantee that the hippocampi of subjects in any particular study will be organizing in that manner. Hence, the data-driven approach to parcellation may frequently be the most accurate and informative approach.

In conclusion, structural differences along the hippocampus have long been thought to underlie meaningful functional differences. While it is true that structure certainly constrains function, structure should not be confused with function— boundaries between structure do not always define boundaries between function. Functional organizations appear to change with experience and directly affect performance on tasks. Nonetheless, there may be an optimal or idealized hippocampal parcellations across tasks. The mICA-10 parcellation map has been seen in both resting state and now also in spatial navigation, and may represent one such optimal organization.

## Materials and Methods

### Participants

Thirty-nine healthy right-handed volunteers were recruited. Five participants were excluded due difficulties with the task and an inability to complete the navigational training. Of the remaining thirty-four participants who completed the study there were 6 males and 28 females (mean age 20.17 years, range 18-24 years). All participants were free of psychiatric and neurological conditions. All participants had normal or corrected-to-normal vision and otherwise met the criteria for participation in fMRI studies. Informed consent was obtained from all participants in accordance with SUNY Geneseo and the University of Rochester’s ethical guidelines. Participants received monetary compensation upon completion of the study.

### Surveys and Demographics

Participants completed a brief survey to gather demographic data and assess their navigational abilities before participant in the study. This survey included a question asking participants about how many hours per week they play video games. It also included both the Navigational Strategies Questionnaire (2) and the Santa Barbara Sense of Direction Scale (24).

### Routes and Navigation Software

Researchers selected the area of Squirrel Hill, PA for this study (see Figure 1A). This region was selected due to the fact that it contained a somewhat grid-like layout of streets, giving subjects a reasonable opportunity to develop spatial maps where they might be able to discover or infer shortcuts between routes, and because it contained a number of visually distinct sub-neighbourhoods, ranging from houses, shops, large fields and golf courses, and even streets with distinctly colored cobblestone. The variation in environments was desired as it was hoped this would help subjects orient themselves as they learned the layout of all of these distinct regions.

A custom software suite was designed in Python to both download images from Google Street View and recreate the Google Street View experience in the lab, to allow participants to virtually navigate and walk through environments. With this software, we could place participants at any position in the neighbourhood and provide them with a panoramic, first person-view of their position. They could use the arrow keys (on non-scanner day) or button box buttons (on scan days) to rotate left or right in 360 degrees. They also had a button to step forward so that they could traverse the environment.

This software was used to download the Squirrel Hill, PA neighbourhood ahead of time and to run the navigation portions of the experiment.

### Navigation Training & Testing

The experiment took place over a seven sessions in a 2-week period (see Figure 1B). Sessions 2, 6, and 7 were scanning sessions, whereas the rest occurred outside the scanner. During the first five sessions, participants were trained to navigate 4 routes (Figure 1A) across 5 training sessions of passive navigation (with the second session being a scanning session, the rest were held outside the scanner; Figure 1B). The four learned routes were relatively non-overlapping but all passed through a central intersection in the middle of town. Each route in the training sessions began and ended at a distinctive landmark and ended and these landmarks were used to generate names for each route. The landmark names were invented by the researchers to be distinctive and informative. The route names were: Grey’s Manor to O’Connor Country Club, Colfax Academy to Cobblestone Way, Red Brick Road to Homewood Estate, CVS Market to the Uptown Lot. At the beginning of each training route these names were presented to participants, along with an image of their destination landmark. During navigation, the name of the destination landmark was present at the top of the screen along with an arrow which directed participants where to go (see Figure 1C). At periodic intersections, the experiment would pause and participants were asked which way they would go next in order to proceed on this route.

Participants could answer left, right, or forward, and were immediately presented with feedback via the arrow updating in the correct direction after their response. Accuracy for these directional quizzes is presented in Figure 1F. Following each training trial a set of recognition tests were presented where subjects where shown scenes from the route they navigated and had to identify whether the scenes came from the route at hand or were being presented in the correct order.

For each subject, the four training routes were randomly assigned to be Routes 1, 2, 3, and 4. In the first session of training, only Routes 1 and 2 were practiced (see Figure 1B). Session 2 was the first scan and all 4 routes were trained during this session. Hence, Routes 1 and 2 were recently experienced whereas Routes 3 and 4 were completely novel, providing us with data on both kinds of routes in this early scan. In Sessions 3, 4, and 5 Routes 1-4 were repeatedly practiced outside the scanner in Sessions 3-5. Each training session took approximately 1.5-2 hours and 11 routes were practiced during each session.

Participants who exhibited steady learning in these conditions were classified into the Strong Learners category and were invited to participate in the sixth experimental session (an active navigation session). Strong Learners showed 80% or greater accuracy on directional quizzes by the 5^th^ training session (see Figure 1F). Participants who exhibited difficulty with the navigational task and who showed poorer learning performance were classified as Weak Learners and not invited to take part in the sixth experimental session. In pilot testing participants who had not learned the routes well floundered during the sixth experimental session, which involved various navigational challenges, and often became frustrated. Given these issues and the fact that we also had budgetary constraints, we allocated the sixth day scans to the Strong Learners only.

The specific trials in the sixth session included Route Following, Route Finding, and Blocked Routes. For Route Following, participants were asked to travel each of the four training routes accurately from memory. For Route Finding, participants were placed at one of the starting landmarks and asked to navigate to the destination landmark from a different route, thus requiring them to find a way to bridge between two known routes. Lastly, for Blocked Routes participants were asked to travel a trained route from memory however, they were informed that the center of town had been blocked off and that when they reached the obstruction they would have to find an alternate route to the destination. We used the experimental software to place artificial road signs around the center square of town and thus physically prevented subjects from traversing those roads, forcing them to find alternate routes to their destination. In total, there were 4 Route Following trials, up to 9 Route Finding trials, and 4 Blocked Route trials.

Participants had a maximum of 8 minutes to navigate any particular route but in practice were not given 8 minutes on every route due to scanner timing limitations—if participants took too long navigating on some trials (or had clearly become lost) the researcher had to skip other trials in order to keep the experiment on track. Thus, due to these complex timing balances, we prioritized collecting all 4 Route Following and all 4 Blocked Route trials, and then also collecting as many Route Finding trials as time would allow.

Finally, on the seventh session participants were presented with each of the four trained routes once, and were guided along the route in the exact same manner as they were in the first five training sessions. This final session was carried out in the scanner so that we could compare passive navigation in Session 2 (early in training) with passive navigation in Session 7 (after training).

### fMRI Image Acquisition & ICA Analyses

Scanning took place at the University of Rochester Center for Advanced Brain Imaging & Neurophysiology. Participants were scanned on a Siemens 3T Prisma MRI scanner. On the first scan day (the second experimental session) a high-resolution 3D MPRAGE T1-weighted pulse sequence image (192 axial slices, 1 mm thick, FOV = 256 mm) was first obtained to register functional maps against brain anatomy. Functional imaging was performed to measure brain activation by means of the blood oxygenation level dependent (BOLD) effect. Functional T2*-weighted images were acquired using echo-planar imagine (84 axial slices, 2 mm thick, TR = 1500 ms, TE = 30 ms, flip angle = 60 degrees, FOV = 200 mm). The native EPI resolution was 128 × 128 with a voxel size of 2mm × 2mm × 2mm.

Images were preprocessed using the FEAT tool in FSL v6.00. Preprocessing involved motion correction (MCFLIRT), brain extraction (BET), spatial smoothing (5mm kernel), high-pass filtering, spatial realignment, and co-registration. Pre-processed data was then transformed into MNI space for the grouped masked independent component analyses (mICA). Hippocampal masks for these analyses were generated from the Harvard-Oxford cortical and subcortical structural atlas by applying a threshold of 50 and binarizing the result. Using these masks, mICA was applied to the functional data using the mICA toolbox v1.18 (6). Replicating the work of Thorp et al. (7), to obtain cluster maps multi-session (group) ICA was selected with the number of components limited to 10 unless specified elsewhere in the Result. To carry out reproducibility analyses, 50 repetitions of split-half reliability sampling was selected in the toolbox, along with the components limited to the dimensionality at hand (often 10 but varied depending on the specific analysis as described in the Results).

Jaccard coefficients were calculated using the Jaccard bash script (https://github.com/KrbAlmryde/Utilities/blob/master/WorkShop/SHELL/jaccard.sh). A custom search algorithm was written in Python to try to maximize the pairings that were used when calculate mean Jaccard coefficients for pairs of cluster maps. Statistical results for all analyses were plotted and analyzed using RStudio v4.2.1

## Acknowledgments

We wish to thank Harris Schwab, Kaitlyn Bertleff, and Luke Bamburoski for help carrying out this experiment. We also wish to thank the staff at the University of Rochester Center for Advanced Brain Imaging & Neurophysiology for their support. This research was supported by NIH (1R15NS104979-01).

